# Studying the effect of conserved tyrosine phosphorylation within SH2 domains

**DOI:** 10.64898/2026.08.13.744714

**Authors:** Gabrielle Martinez, Mindolay Fike, Madhuri Sosale, Swathi Shekharan, Kristen M. Naegle

**Affiliations:** Department of Biomedical Engineering and the Department of Genome Sciences, University of Virginia, Charlottesville, Virginia, United States of America

**Keywords:** Signaling, Protein, Post-Translational Modification, Phosphorylation

## Abstract

SH2 domains are phosphotyrosine-binding modules that play a critical role in cell signaling by mediating protein-protein interactions. While tyrosine phosphorylation has been shown to impact SH2 domain function in signaling, the specific effects of phosphorylation at different sites within the domain remain poorly understood. In this study, we selected two conserved regions of tyrosine phosphorylation within SH2 domains, near conserved binding interface residues, and developed approaches to evaluate the impact of those sites on ligand binding. Using a modified dot blot assay to screen phosphomimic mutations, we studied specific tyrosine residues within the PTPN11-N, LYN, and SYK-C SH2 domains, finding that the PTPN11 N-terminal site (Y63) modulates the specificity, reducing binding of physiologically relevant substrates. Our findings provide new insights into the regulatory mechanisms governing SH2 domain function and highlight the importance of site-specific phosphorylation in modulating protein-protein interactions in cell signaling pathways.

## Introduction

Src Homology 2 (SH2) domains are responsible for binding tyrosine phosphorylated proteins and are found in a wide variety of signaling proteins, including kinases, phosphatases, and adaptor proteins. In the human proteome, there are 120 SH2 domains across 110 proteins (1, 2). They are approximately 100 amino acids long and share a conserved fold that consists of a central *ω*-sheet flanked by *ε*-helices and are best known for recognizing phosphotyrosine (pTyr) containing peptides (3, 4). Through these interactions, SH2 domains play a central role in phosphotyrosine-mediated signaling networks. Their importance is so high, that it was proposed that their emergence was essential to the evolution of tyrosine kinase signaling (5).

Despite sharing a highly conserved structure, SH2 domains exhibit distinct specificity in their binding interactions. Binding occurs via a conserved arginine residue, which coordinates pTyr engagement, while residues surrounding the pTyr contribute to specificity (4, 6). As a result, SH2 domains can recognize distinct signaling partners based on sequence motifs typically spanning positions +2 to −4 relative to the pTyr (7, 8). Historically, SH2 domain specificity has been characterized using peptide library screens, structural studies, and quantitative binding assays, which have revealed that individual SH2 domains possess unique preferences for residues surrounding the phosphotyrosine. These selective interactions are critical for determining which signaling complexes form downstream of receptor tyrosine kinase (RTK) activation and therefore help direct specific cellular outcomes. However, understanding SH2 domain specificity remains challenging because these interactions are often low to moderate affinity and are influenced by both local sequence context and the broader cellular environment (9). Because of this, predicting SH2 domain binding partners remains a complex problem.

As the field still seeks to fully understand specific SH2 domain interactions based on how the pocket creates specificity for partner targets, another complexity has also emerged. Specifically, SH2 domains themselves are heavily tyrosine phosphorylated, with over 190 known pTyr sites across the 120 SH2 domains in the human proteome (10). A few limited studies have demonstrated a few of these SH2 domain modifications can have significant consequences on SH2 domain binding and selectivity. For example, Stover et al. demonstrated that SRC Y213 phosphorylation reduced binding with the SRC C-terminal tail negative regulatory sequence, but without altering binding to a ligand of the EGFR C-terminal tail (11). Similarly, Couture et al. found phosphorylation of LCK Y192 reduced binding recruitment in signaling and corresponded reduced overall signaling response to T cell activation (12). More recently, Jin et al. found LYN Y194 appeared to generally reduce binding to a wide variety of partners, though there may be some indication this was selective (13). These findings indicate that phosphorylation has the potential to regulate SH2 domains, altering their engagement in signaling networks on the fly. However, prior studies are currently limited to one structurally conserved site, at the end of the *ω*-E strand, within several members of closely related homologs (in the SRC family kinases). As a result, it remains unclear how widespread phosphorylation-dependent SH2 regulation is, which structural features distinguish regulatory sites from non-regulatory sites, and how these modifications influence phosphotyrosine signaling networks. Addressing these questions is essential for understanding how SH2 domains dynamically regulate cellular communication.

To begin addressing these knowledge gaps, we identified multiple positions of conserved phosphorylation across the SH2 domain family and, using structural mapping of SH2-ligand interfaces using CoDIAC, we identified sites likely to affect ligand interactions, given their proximity to ligand binding positions (10). One of these conserved positions is the well-characterized phosphotyrosine site found in several SRC family kinases, which corresponds with locations that interact with the specificity region of the binding pocket.

Additionally, another highly conserved position, is located closer to the pTyr coordination region. We therefore hypothesized that phosphorylation at conserved positions near the ligand-binding interface would alter SH2-domain phosphopeptide recognition (those that impact closest to the pTyr interaction region) and binding specificity (those that correspond with the specificity region). To test these hypotheses, we selected representative SH2 domains containing phosphotyrosine sites within the two conserved regions of the ligand-binding interface. Because stable site-specific phosphorylation of SH2 domains remains experimentally challenging, phosphomimic mutations were used to approximate the introduction of negative charge associated with phosphorylation. We then developed and applied a dot blot overlay assay to compare the phosphopeptide binding profiles of phosphomimic and wild-type SH2 domains across diverse peptide substrates. We found variable results in the ability of phosphomimic mutations to produce effects on SH2 domain binding, with several showing no effect. However, we found substantial effects for PTPN11 Y63, where only the acidic amino acid mutation resulted in a complete loss of recognition for physiologically relevant ligands (GAB1 and PD-1), while retaining binding with other ligands.

## Results

To hypothesize functional effects of the 190 tyrosine phosphorylation sites in SH2 domains, we used structural analysis to identify sites of conserved tyrosine phosphorylation and their relationship to SH2 domain ligand binding sites (10). Analysis of SH2 domain structural alignments revealed two regions of interest containing recurrent phosphotyrosine sites: one located near the phosphotyrosine-binding pocket and a second located within the specificity-determining region of the domain (Fig. 1). Representative SH2 domains containing phosphosites in these regions were selected for experimental characterization. To evaluate the functional consequences of phosphorylation, we next developed an inexpensive and scalable phosphopeptide-binding assay capable of detecting changes in SH2-domain binding behavior. This assay was then used to compare wild-type and phosphomimic mutants and assess the potential regulatory effects of phosphorylation at conserved SH2 domain positions.

**Fig. 1.**
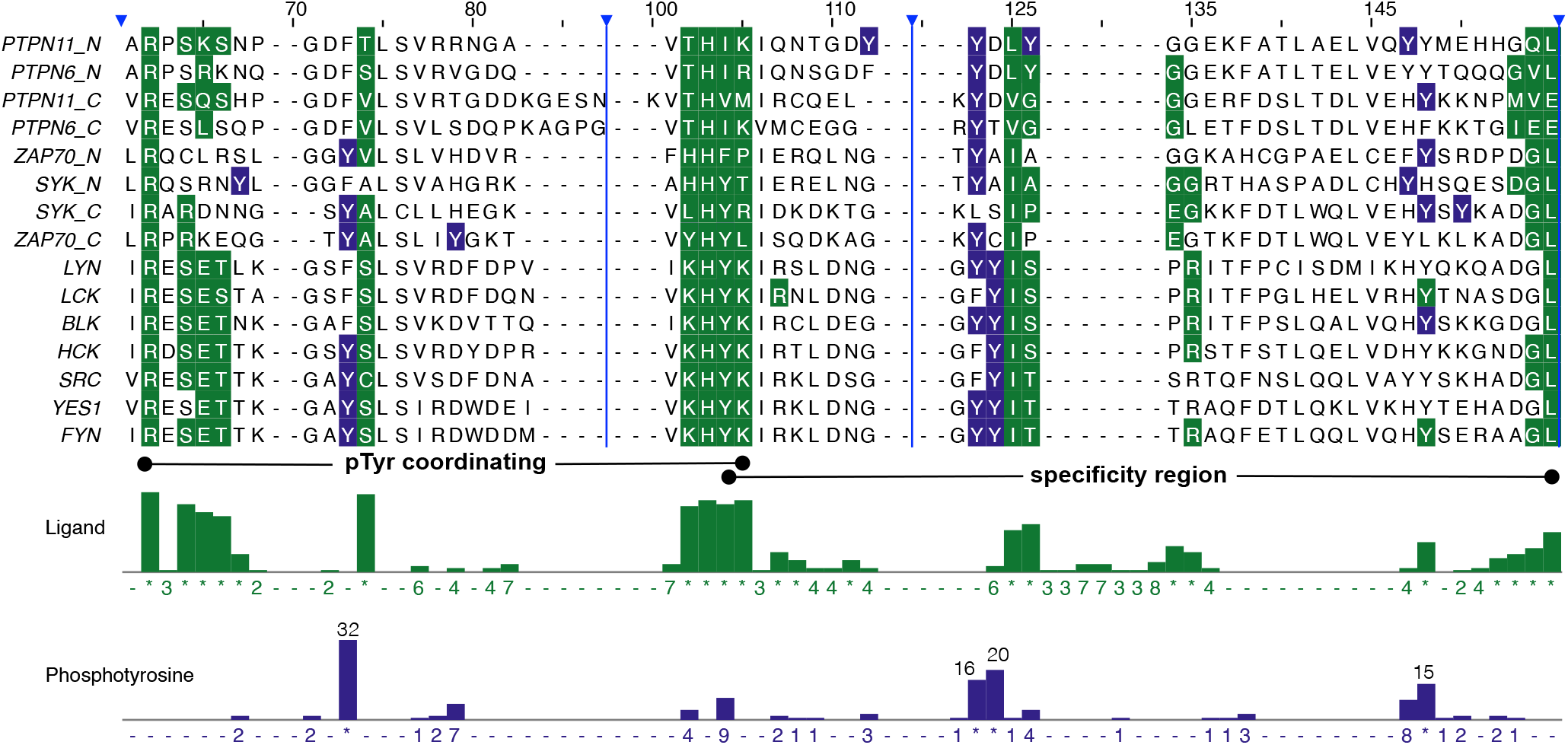
The global structural alignment of the reference from Kandoor et al. (10) is shown with residues that interact with ligands (green) and documented phosphotyrosines (blue). A subset of proteins and alignment positions are shown for reference. Total number of domains with contacts or pY at a position are given on the bottom tracks and prior work has shown that the pTyr coordinating region occurs in the first N-terminal half and the specificity binding region in the C-terminal half (10). Positions selected for study here are alignment column 73 and 123-124.

### Selection of SH2 domains and phosphotyrosine sites for testing

Phosphorylation of SH2 domains is extensive, with almost 200 phosphorylated tyrosines identified across the 120 human SH2 domains. We focused on the combination of conservation – conserved presence of a phosphorylated tyrosine in a conserved structural position of the SH2 domain to prioritize sites for study. Additionally, we used their relationship to the ligand binding interfaces to hypothesize effects and prioritize conserved sites with likely impacts on ligand binding for further study. We developed a comprehensive domain-based structural extraction approach, CoDIAC, which we had previously applied to SH2 domains. CoDIAC extracts structural information for all available SH2-containing structures, mapping all residues in the SH2 domain that are within 4Å of a ligand interface. The SH2 domain binding interface consists of several functionally distinct regions, including the canonical phosphotyrosine-binding pocket, which engages with the ligand pTyr residue, and specificity-determining regions that engage residues C-terminal to the phosphotyrosine to define specificity. Mapping conserved phosphorylation sites onto these regions revealed two key regions of highly conserved tyrosine phosphorylation (Fig. 1) – alignment position 73 (in the pTyr coordinating region) and alignment positions 123/124 in the specificity region, specifically positions found to coordinate the +3 binding region of the ligand (10). A third region of conserved pY exists at the N-terminal end of SH2 domains (alignment position 22), but this region is not near the ligand binding area and based on prior results, may be more likely to occur at intraprotein interfaces in multidomain proteins (10). Finally, a fourth region also exists (alignment positions 146-147), where a relatively large number of pY exist and overlap with some interaction residues. However, this interaction interface is more variable, belonging to only certain families, and so it is likely less universally conserved across SH2 domains. Overall, conservation analysis suggests that there may be two key points of regulation for tyrosine phosphorylation on the interface of SH2 domain binding shared by most SH2 domains – on a position that may impact overall recognition (being in the pY binding region) and positions that may affect specificity (being in the +3 determinant region).

The two positions that appear near globally conserved ligand binding residues account for 68 pY sites in SH2 domains, or approximately 36% of all pY sites recorded to date. In the phosphotyrosine-binding region (alignment position 73), there are 32 pY sites currently annotated with an additional 29 domains that contain a tyrosine residue at this position (i.e. half of SH2 domains have a tyrosine position in this location and half of those have been observed as phosphorylated). Similarly, at the specificity-determining region (alignment positions 123/124) there are 53 tyrosines, where 36 have been reported as phosphorylated. Notably, phenylalanine is also commonly observed at both positions, accounting for 34 sites at position 73 and 21 total sites in positions 123/124, suggesting evolutionary conservation of an aromatic residue while permitting variation in the ability to be phosphorylated. These observations support the hypothesis that phosphorylation at specific positions within the SH2 domain may represent a conserved regulatory mechanism, rather than isolated events.

Having identified specific positions of interest, we next wished to focus on specific representative SH2 domains for further study. From this analysis, we selected three representative SH2 domains: PTPN11 N-SH2, LYN SH2, and SYK C-SH2. These domains were chosen, because they contain conserved phosphorylation sites positioned within structurally important binding regions, while also representing diverse signaling protein families and functions. PTPN11 N-SH2 and LYN SH2 were of particular interest because both contain phosphotyrosine sites located near the specificity-determining region of the binding interface, yet the sites oc-cupy slightly different structural positions (Fig. 1). The LYN SH2 domain also allowed us to test the accuracy of a phosphomimic mutation at Y194, as phosphorylation of Y194 has previously been seen to reduce binding to known binding partners (13). In contrast, the neighboring PTPN11 Y62 and Y63 sites have been implicated in regulation of PTPN11 activity, but have not been extensively studied with respect to their effects on SH2-mediated ligand recognition. SYK CSH2 was selected to represent a separate region of phosphosites located adjacent to the phosphotyrosine-coordinating pocket rather than the specificity determining region. This allowed us to test a separate hypothesis generated from the CoDIAC analysis: that phosphorylation near residues directly involved in phosphotyrosine recognition would be more likely to alter overall ligand binding, where phosphorylation near specificity determining residues would influence ligand selectivity. Finally, these specific SH2 domains were of interest as they covered multiple tyrosines in the region, where family members lack phosphorylation ability – PTPN11 Y62 homology is a phenylalanine in PTPN6 and several SRC family kinases lack tyrosine at LYN Y193 location.

### Assay development for monitoring SH2 domain specificity

To evaluate how phosphorylation within SH2 domains influences ligand binding, we first aimed to establish a robust method for monitoring SH2-ligand interactions. Far westerns, a commonly used method for studying protein-protein interactions, was initially considered because it detects interactions between SH2 domains and substrates using a relatively simple protocol (14, 15). As an initial test, we evaluated whether endogenous phosphotyrosine-containing proteins in mammalian cells could allow for the detection of distinct binding patterns for SH2 domain overlay assays. Lysates prepared from U251 cells treated with pervanadate were analyzed by far-western blotting after incubation with purified GST-fusion SH2 domains. Although phosphotyrosine-containing proteins could be detected, the resulting overlay signal was very weak, even with the advantage of dimerization presented by GST fusion (15), and nonspecific GST antibody binding made it difficult to distinguish domain-specific binding interactions from background noise from GST antibody alone (Fig. S1). These observations suggested that a higher density pTyr source would be necessary for semiquantitative binding measurements.

To address this limitation, we utilized SISA-KiT, a synthetic kinase toolkit to produce recombinant phosphotyrosine-containing substrates, which involves an SH3-fused kinase targeted to a polyproline-fused substrates of interest (16). We co-expressed two substrates known to bind the N-terminal PTPN11 SH2 domain, PD-1 pY248 and GAB1 pY627,which we had found in other work can be sufficiently phosphorylated as reagents for antibodies (16) and SH2 domain binding (17). When tested in the far-western format, these phosphoproteins generated substantially stronger binding signals than those observed with mammalian lysates, while maintaining low background (Fig. S1). The resulting increase in detection of SH2-mediated interactions demonstrated the feasibility of using recombinant phosphoproteins as substrates for binding assays, while being significantly less expensive than synthetic peptides to produce. In addition, we evaluated whether substrate purification was necessary. Comparison of purified PD1 pY248 and crude GAB1 pY627 lysate revealed that both preparations were sufficient for SH2 domain binding (Fig. S1). Hence, it became clear that relatively low expense of cloning and co-expression using this approach could be used to produce high signal to noise approach in SH2 domain binding assays.

As this was now a more targeted approach to assessing binding changes, we transitioned from a traditional far western format to a dot blot overlay assay. Rather than separating the lysates by SDS-PAGE, individual phosphoprotein substrates are spotted directly onto membranes and probed with purified SH2 domains, allowing for increased density and higher throughput. To ensure that the binding observed was specific to the substrates, a tyrosine-free control was included for each kinase used to phosphorylate target peptides. This control accounted for off-target phosphorylation of endogenous bacterial proteins, which could contribute to non-specific SH2 domain binding. This negative control establishes whether SH2 domain binding is specific to the target and we found across all controls that background binding to E. coli lysates expressing tyrosine kinases is minimal (Fig. S4).

Having established the key components of the assay, we expanded the system to generate a collection of phosphopeptide substrates representing known SH2 domain ligands. To maximize experimental throughput, four candidate substrates were selected for each SH2 domain, based on available data. We cloned the target sequences into an expression vector and tested phosphotyrosine yield based on phosphotyrosine antibody binding, selecting a kinase partner that produced viable substrate phosphorylation (ultimately using ABL and SRC kinases for the panel of 11 phosphopeptides). In order to improve assay interpretability, we screened each domain against a serial dilution of batch phosphopeptide stocks (Fig. 2) to identify a 4 dilution range that was linear for each domain.

**Fig. 2.**
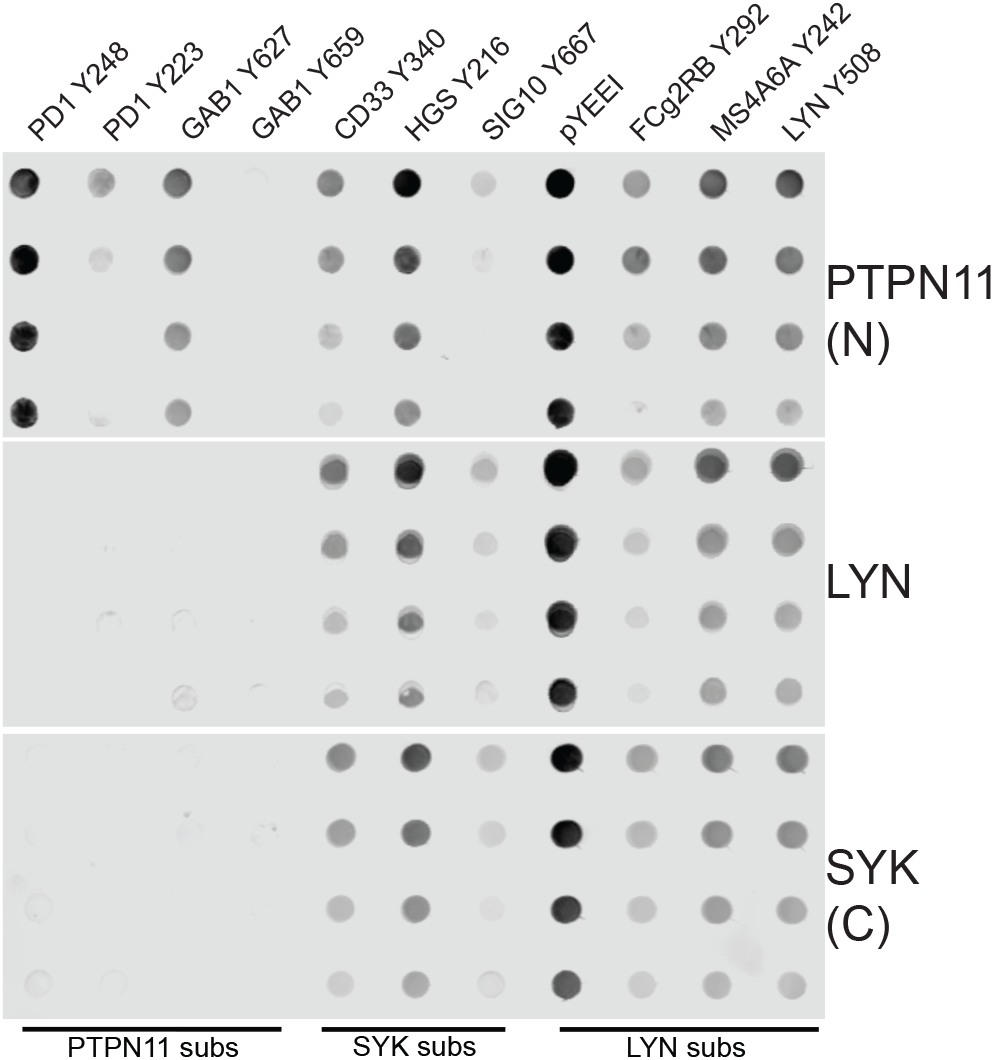
Full panel phosphopeptide screens using wild-type PTPN11 N-SH2 (top), LYN SH2 (middle), and SYK C-SH2 (bottom).

This screen highlights innate SH2 specificity differences between our proteins, but also some interesting overlaps in substrate partners. For example, only PTPN11 N-SH2 binds to the PD1 and GAB1 substrates, though binding to the GAB1 pY659 was poor, but all three domains strongly bind the synthetic YEEI peptide that is the idealized partner for LYN substrate binding.

Notable interactions identified during the initial screens were subsequently evaluated using replicate dot blot assays. For these experiments, the four strongest binders were applied in triplicate using adjusted protein concentrations on a single membrane to assess technical reproducibility within an experiment (Data S2). The fourth quadrant was dedicated to the application of negative and positive controls (Fig. S4, Fig. S5, Fig. S6). Although minor differences in overall signal intensity were observed between experiments, the relative binding patterns of the wild-type SH2 domains remained highly consistent across replicate membranes. This reproducibility demonstrated that the assay reliably captured SH2 domain binding preferences and provided a robust platform for comparing phosphomimic and wild-type domains.

### Using phosphomimic mutagenesis to test multi-sub-strate binding effects

Having established a reproducible and easy approach to profiling selectivity, we next wished to test the specific hypotheses regarding the role of phosphorylation within the SH2 domains at the conserved sites identified. Unfortunately, producing phosphorylation of the SH2 domains is highly challenging. There are methods for synthetic amino acid incorporation, but they are expensive and produce low yields (18) and have complex conditions (for example, deprotecting a neutralizing group by placing folded protein at pH 1.0 for 16 hours (19)). The advantage of the toolkit we used to produce substrate is that it produces high yields of native tyrosine phosphorylation. However, it is non-specific and so it requires mutagenesis of possible off-target tyrosines. Additionally, GST fusions cause significant challenges (presenting a sink for kinases) (16). We tested whether we could replace the GST with another detection approach – using an ALFA-tag, which is tyrosine free, N-terminal fusion and an IR-dye conjugated nanobody with mutations of the tyrosine without interest. However, the GST dimer is responsible for helping amplify signal from the relatively low affinity interactions (15, 20), which are further impacted by multisite non-phosphorylatable mutations at off-target tyrosines (Fig. S3), and so these approaches were not sufficiently robust as a method for measuring SH2 domain interactions. Instead, we used mutagenesis to glutamic acid (E) to mimic the negative charge of a modified tyrosine in the SH2 domain at the identified sites of interest. Phenylalanine (F) mutations were also introduced in the same position as the phosphomimic mutations as a control to confirm whether the mutation of the tyrosine alone is leading to binding changes, or if it is due to the presence of a negatively charged amino acid. Here, we will discuss the study of Y62 and Y63 in PTPN11 N-terminal SH2 domain, Y203 of the SYK C-terminal domain (along with S202, a serine phosphorylation site), and Y193 and Y194 of the LYN SH2 domain using mutagenesis, com-bined with the dot blot overlay assays designed to screen for specificity changes.

### PTPN11 N-SH2 domain binding effects of Y62 and Y63

PTPN11 (SHP2) is a phosphatase containing tandem SH2 domains that functions as a key mediator of receptor tyrosine kinase (RTK) signaling in the RAS/MAPK pathway (21, 22), and studies have shown that it is necessary for survival of RTK-driven cancer models (23). PTPN11 is activated through the binding of its N-SH2 domain to phosphotyrosine-containing ligands, which induces a conformational change – releasing the autoinhibitory interaction between the N-SH2 and the phosphatase domain and activating the phosphatase (24). Recent studies have demonstrated that phosphorylation of the N-SH2 domain at Y62 leads to increased PTPN11 activity due to disruptions at the N-SH2:PTP interface preventing PTPN11 from returning to an inactive state, and this has been seen to have negative consequences in AML, where pY62 led to the resistance to allosteric inhibitors (25–27).The neighboring Y63 site has also been suspected to stabilize the active conformation, but has not been studied nearly as much, motivating our interest in testing the effects of phosphorylation at this site as well. CoDIAC-based structure analysis is consistent with the idea that Y62 is important at the SH2:PTP interface and highlights that Y63 is closer to conserved ligand binding interface residues.

We isolated each site by mutagenesis (Y62E and Y63E, along with Y62F and Y63F control mutations). We compared binding of these mutants, compared to wild type (WT) domain binding on four phosphopeptide substrates that showed dynamic binding range to WT PTPN11 N-SH2 domain (Fig. 2): PD1 pY248, PD1 pY223, GAB1 pY627, and HGS pY216.The Y62E mutant did not display any significant changes to binding to the tested substrates. Although binding to PD1 pY223 appeared to decrease, a similar decrease was also observed in the Y62F mutant, suggesting that this change in specificity was due to the mutation of the tyrosine alone, not the introduction of a negative charge (Fig. 3). In contrast, the Y63E mutation displayed a distinct binding profile. Binding was significantly reduced for three of the four substrates, while interaction with HGS pY216 remained intact (Fig. 3). This selective change in substrate recognition may be due to the proximity of Y63 to the specificity determining region of the SH2 domain, as identified by CoDIAC analysis. Interestingly, the site in PTPN11 adjacent to Y63 (same conserved position as LYN 194) is one of the only instances of a negatively charged amino acid present in this position within all SH2 domains, suggesting that phosphorylation at this position could induce more significant structural changes that alter the preferred binding motif, resulting in changes in substrate specificity rather than a loss of binding.

**Fig. 3.**
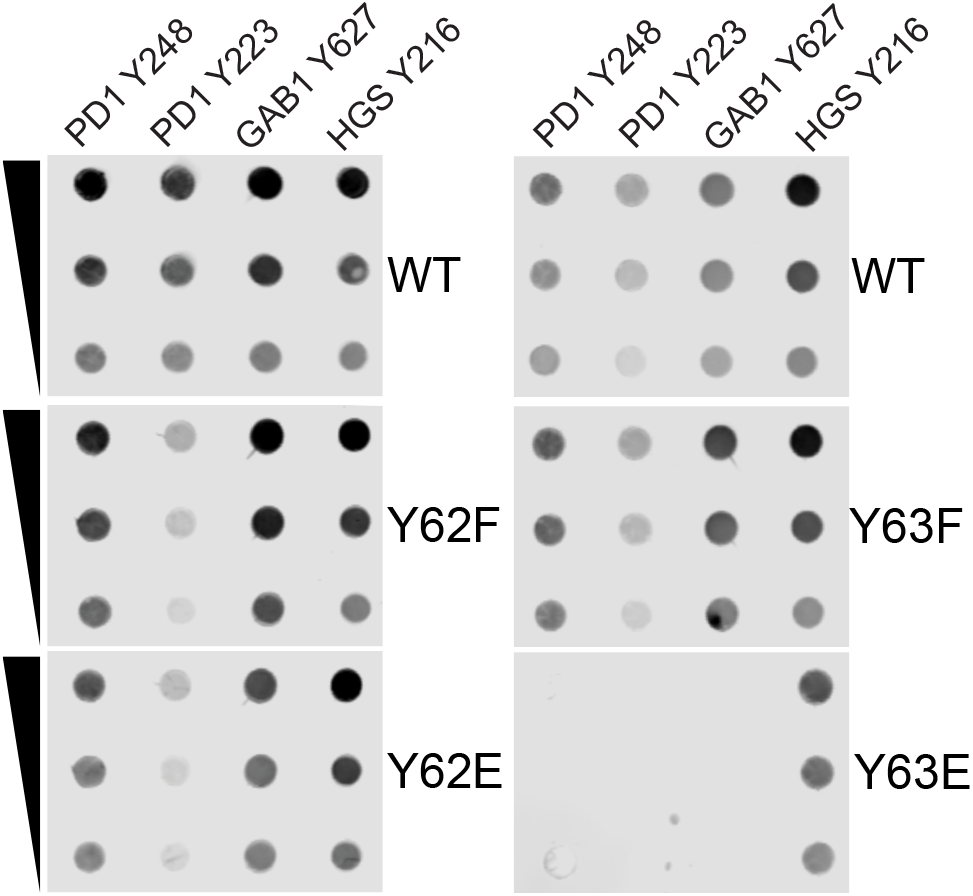
PTPN11 N-SH2 binding of Y62 and Y63 phosphomimics and controls. Dot blot overlay assays were performed with equal concentration of PTPN11 N-SH2 domain recombinant proteins against the same 2-fold serial dilutions of substrate lysates. Left panels Y62 mutants: including the WT replicate performed with Y62E and Y62F. Right panels Y63 mutants: including the WT replicated performed with the Y63E and Y63F.

Given the selectivity switch for the Y63E mutant, we wished to understand other possible binding changes and subsequently screened Y63E against the total panel of 11 substrates. This expanded analysis revealed that the Y63E mutation selectively lost binding to the PD1 and GAB1 phosphopeptides, while binding to the remaining substrates was preserved. Interestingly, the loss in binding was restricted to the phosphopeptides selected based on their relevancy to known binding interactions with PTPN11 N-SH2 domain (Fig. S1). Together, these results suggest that a negative charge at Y63 alters the selectivity of the PTPN11 N-SH2 domain to physiologically relevant substrates, rather than reducing its overall binding capacity.

### LYN SH2 domain binding effects of Y193 and Y194

LYN is a member of the Src family kinases, and plays important roles in both immunoreceptor tyrosine-based activation motif (ITAM) and immuoreceptor tyrosine-based inhibitory mofit (ITIM) signaling pathways (28, 29). SH2 domains within the SFK family all contain a highly conserved phosphotyrosine site corresponding to LYN Y194 that have been seen to be phosphorylated in many different types of cancers, including AML and NSCLC. Structurally, this residue is located within the specificity determining region, located close to amino acids interacting with the +3 position on the ligand, suggesting that phosphorylation in this area could affect the binding specificity and affinity for ligands. This is supported by a study previously done on LYN Y194, where it was found that phosphorylation of this site led to decreased binding to known binding partners (13). In addition to Y194, LYN contains a neighboring phosphotyrosine site that is conserved only within a subset of SFKs and has not been functionally characterized, and possibly complicated interpretation of the prior study, which did not explicitly control for alternate pY sites in the LYN SH2 domain (13). This offered an opportunity to evaluate whether distinct effects could be detected between the adjacent positions.

To evaluate the effect of phosphorylation at these sites, we generated phosphomimic (Y->E) and control (Y->F) mutants for both Y193 and Y194 and tested their binding to four phosphopeptide substrates: pYEEI, FcgRIIb pY292, MS4A6A pY242, and HGS pY216. Three of these substrates were selected based on the binding assays done by Jin et. al (13) for LYN pY194, while HGS pY216 was included because it was found to be a strong binder in our initial screen (Fig. 2). In contrast to our hypothesis, neither the Y193E nor Y194E mutant exhibited significant changes in binding relative to the WT domain (Fig. 4). Given the previously reported effects of Y194 phosphorylation on LYN binding interactions, these results suggest that the phosphomimic mutation may not fully capture the charge and structural changes caused by phosphorylation at these sites. Since Y193 and Y194 are adjacent phosphotyrosine sites, we also generated a double mutant (Y193E/Y194E) to test whether simultaneous modification of both residues might produce effects that were not apparent in the individual mutants. However, similar to the individual mutants, the double mutant failed to produced detectable changes in binding to any of the tested substrates (Fig. S5).

**Fig. 4.**
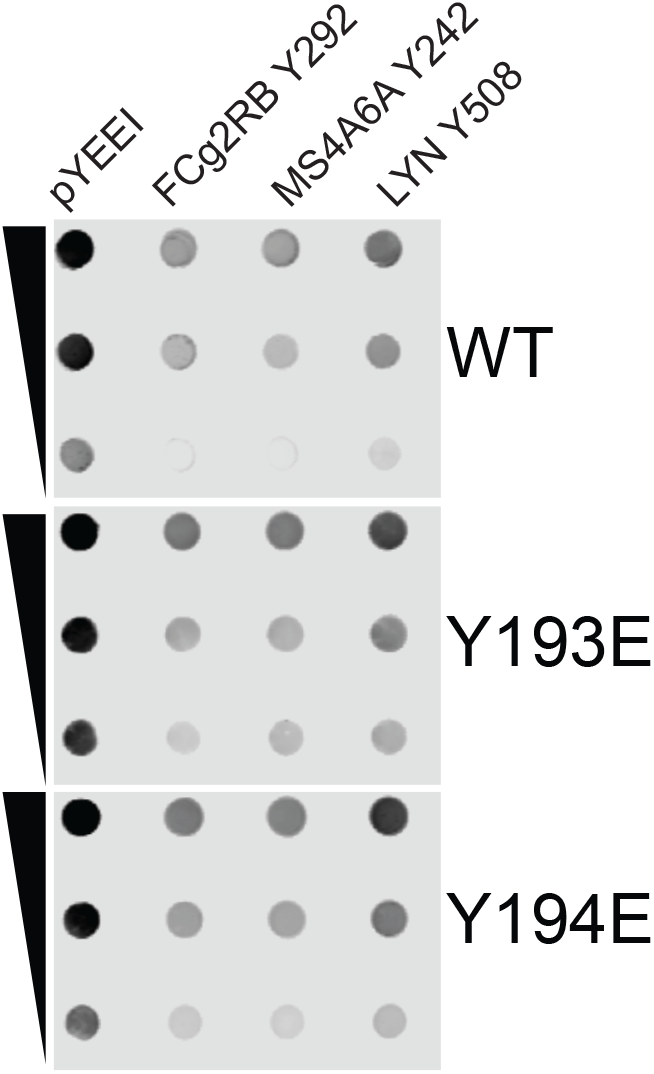
LYN SH2 binding of Y193 and Y194 phosphomimics. Dot blot overlay assays were performed with equal concentration of LYN SH2 domain recombinant proteins against the same 2-fold serial dilutions of substrate lysates.

### SYK C-SH2 domain effects of S202 and Y203

The SYK C-SH2 domain was chosen because it contains a phosphorylation site (Y203) that is in the conserved region that coordinates directly the ligand pTyr. Hence, we posited this position would be more likely to disrupt all binding, rather than change specificity alone. Additionally, SYK C-SH2 domain represented an interesting opportunity to also consider adjacent phosphoserine (S202) that is also a site of conserved phosphorylation. Similar pS/pY pairings have been observed in other SH2 domains in this region, prompting us to investigate whether phosphorylation at S202 could provide additional insight into how phosphorylation near the binding interface influences SH2 domain function.

To test this hypothesis, we generated phosphomimic mutations at both positions (S202E and Y203E) and assessed binding to the four highest affinity substrates identified in the SYK C-SH2 WT peptide screen: CD33 pY340, HGS pY216, FcgRIIb pY292, and MS4A6A pY242. Surprisingly, neither the S202E nor Y203E mutation produced significant changes in binding in either the quadrant screens (Fig. 5) or the full panel screen (Fig. S9). These results suggest that, if there is an effect of phosphorylation at these sites, that the phosphomimic is not capable of capturing these differences. We anticipate that phosphomimics are more suitable for capturing the size and charge of serine than it is of tyrosine. Hence, S202E might better reflect that phosphorylation at this site does not significantly impact binding, at least as measured using this approach.

**Fig. 5.**
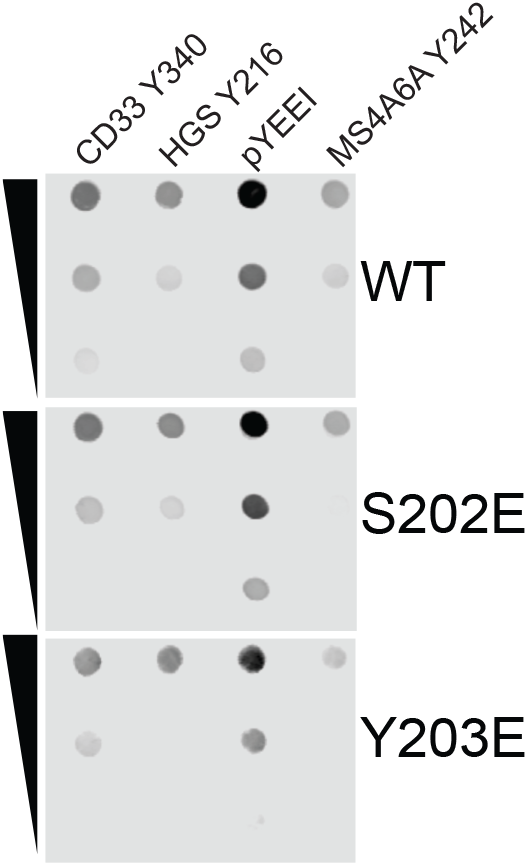
SYK C-SH2 binding of S202 and Y203 phosphomimics. Dot blot overlay assays were performed with equal concentration of SYK C-SH2 domain recombinant proteins against 2-fold serial dilutions of phosphorylated CD33 Y340, HGS Y216, YEEI, and MS4A6A Y242.

## Discussion

This work sought to explore the possibility that conserved regulation of the SH2 domain occurs by tyrosine phosphorylation, expanding beyond the LYN Y194 site that has been previously studied in SRC family kinases. In this work, we also developed fast and inexpensive approaches to developing ligand partners for SH2 domains and reproducible assays that helped us explore specificity across several substrates at once. Unfortunately, the relatively low affinity of SH2 domain binding still requires experimental tricks for boosting signal, such as through GST-based dimerization. This ultimately limited our ability to study the native phosphorylation of SH2 domain, instead using phosphomimics to introduce the negative charge within the SH2 domain. These mim-ics are more likely to introduce a false negative result than a false positive – i.e. a failure to see a change may not be the case in a natively phosphorylated domain, whereas if the negative charge specifically alters binding, compared to the phenylalanine control, than it a native pY is likely to have that effect or more severe. For example, whereas Jin et al. (13)phosphorylated LYN domain with a recombinant kinase and observed alterations in binding, our Y194E mutant did not show substantial changes to binding. However, LYN has three total tyrosines, all of which are phosphorylated, and the mass spectrometry based analysis performed in the Jin study would have limitations in establishing the stoichiometry of these sites as well. Hence, we cannot rule out possibilities that SH2 pull down differences in phosphopeptides from pervanadate treated lysates in that study was partly attributable to complex mixtures of LYN phosphospecies, nor can we assume that Y194E mutation is representative of pY194 SH2 domain function. Additionally, although Y63E of PTPN11 demonstrated significant specificity changes, it is possible those are stronger changes or more global across partners with the larger, bulkier, and more charged phosphorylated tyrosine. Hence, further directed study of SH2 domain phosphorylation may require integrated approaches across many types of experiments to fully understand the consequences of SH2 domain phosphorylation.

The strongest evidence for phosphorylation-dependent modulation of ligand recognition, in this study, was observed within the PTPN11 N-SH2 domain. Previous studies have demonstrated that phosphorylation of Y62 promotes PTPN11 activity by stabilizing the active conformation of the protein, with some suggestion that Y63 can support this as well (27). Our results suggest that the Y63 residue may serve an additional regulatory role in ligand selectivity. Unlike Y62, Y63 is positioned at the beginning of a *ω*-strand that contains a portion of the specificity determining region of the binding interface. The observation that mutation at Y63 altered binding to selected substrates raises the possibility that phosphorylation at this site influences not only PTPN11 activation state, but also the substrates that can engage the N-SH2 domain. This mechanism could have important implications for PTPN11 activity and signaling – engagement of the N-SH2 domain by pTyr-containing ligands is required for activation of phosphatase activity, and many studies have suggested that the N-SH2 domain contributes more strongly to phosphopeptide binding than the C-SH2 domain across a range of substrates (30, 31). If phosphorylation of Y63 alters the specificity of the N-SH2 domain, this modification would regulate not only enzyme activity, but also the specific recruitment of the phosphatase to substrates via alterations in SH2 domain selectivity. The close proximity of Y62 and Y63 highlights the possibility that neighboring pY sites may coordinate different aspects of regulation.

The observed changes in binding specificity for the PTPN11 N-SH2 domain may have important implications for PTPN11-mediated signaling. PTPN11 is recruited to numerous phosphorylated signaling proteins through its tandem SH2 domains, where it functions as a critical reg-ulator of both immune and growth factor signaling pathways. One well-characterized interaction is with the immune checkpoint receptor PD-1, which contains two phosphorylated motifs within its cytoplasmic tail: an immunoreceptor tyrosine-based inhibitory motif (ITIM) centered on Y223 and an immunoreceptor tyrosine-based switch motif (ITSM) centered on Y248 (32), both of which were used as substrates in the binding assays. Studies have demonstrated that the N-SH2 domain preferentially engages the phosphorylated Y248 with high affinity (33, 34), while binding with Y223 is much weaker. Simultaneous engagement of Y248 by PTPN11 N-SH2 and C-SH2 stabilizes PTPN11 recruitment and relieves N-SH2-mediated autoinhibition, resulting in phosphatase activation (34). Consequently, phosphorylation-dependent changes in N-SH2 domain specificity could alter the efficiency of PTPN11 recruitment to PD-1, potentially affecting the magnitude or duration of inhibitory signaling in T cells. In a dynamic study of TCR activation by Chylek et al. (35), the conserved site on PTPN6, Y63, was immediately and strongly phosphorylated upon response to activation (increasing 2-fold within 30-seconds and 4.5-fold within 60-seconds of activation). Like PD-1 recruitment, GAB1 also recruits PTPN11 via strong bivalent interaction with Y627 and Y659 (36), reaching 6nM interaction affinity (17), and promoting downstream ERK signaling (37). Hence, it seems likely that PTPN6 and PTPN11 Y63 phosphorylation might serve as a mechanism to actively break the very strong interactions driven by bivalency.

Beyond functional results, the conservation analysis provides some evolutionary insights into SH2 domains. Several phosphotyrosine sites identified in this study exhibit patterns of conservation and divergence across proteins in the same family. For example, Y63 is conserved in both PTPN11 and PTPN6, whereas Y62 is only present in PTPN11. In LYN, the pTyr site corresponding to Y194 is broadly conserved across SRC family kinases, while neighboring tyrosines (equivalent Y193) are not. In some SFKs, this position is replaced by phenylalanine, preserving a similar structure while eliminating the ability to be phosphorylated. These substitutions suggest that evolution may have not only preserved phosphotyrosine sites, but selectively removed them when phosphorylation-dependent regulation is not advantageous. Interestingly, the Y62 site of PTPN11, which appears to be important to catalytic regulation is phenylalanine in PTPN6, whereas they both conserve Y63 and both are observed to be phosphorylated wide variety of tissues and conditions (38).

Our findings support the concept that SH2 domains can be regulated by phosphorylation and that this is important to shifting the paradigm of understanding in cell signaling. SH2 domains are traditionally viewed as fixed readers of phosphotyrosine signaling, functioning downstream of kinase activity to recruit proteins into signaling complexes whose partnerships are driven entirely by the regulation of the ligand. However, the presence of conserved phosphorylation sites within ligand binding interfaces suggests that SH2 domains may also serve as targets of regulatory phosphorylation. This mechanism would provide a way to dynamically tune signal-ing networks by altering the affinity or specificity of phosphotyrosine recognition. In this framework, phosphorylation would not simply serve as binding sites for SH2 domains, but could also modify the rules by which the interactions occur. Possibly also captured by the conserved analysis is that the SH2 domain might serve as a scaffold on which downstream serine/threonine kinases can feedback to regulate SH2 domain proteins. Although the SYK S202 site we studied here did not show binding changes for the substrates tested, it is possible that other sites of conserved serine/threonine phosphorylation, such as directly on residues that interact with the ligand pTyr are capable of that, and further studies to understand this possible feedback regulation may be a key to understanding overall signaling dynamics and control.

## Materials and Methods

### Generation of Recombinant SH2 domains

Wild type and mutant SH2 domains were produced in pGEX vectors containing an N-terminal GST tag and C-terminal 6xHis tag. Site-directed mutagenesis was performed on wildtype SH2 domains using the Agilent QuikChange Lightning Site-Directed Mutagenesis Kit (Catalog #210518) After the mutated plasmids were transformed into DH5*ε* cells, Sanger sequencing was performed on the miniprepped DNA (QIAprep Spin Miniprep Kit, Catalog #27106) to validate the sequences. Sequences are listed in Data File 1. The sources for the original SH2 domains were: PTPN11: from pBABE-PTPN11 which was a gift from Dr. Matt Lazzara (39). SYK: pDONR223-SYK was a gift from William Hahn and David Root (Addgene plasmid #23907; http://n2t.net/addgene:23907; RRID:Addgene 23907)(40). LYN: pDONR223-LYN was a gift from William Hahn and David Root (Addgene plasmid #23905; http://n2t.net/addgene:23905; RRID:Addgene#23905) (40).

### Generation of Recombinant Phosphopeptide Substrates

Peptide substrates for LYN and SYK were codon optimized for E. coli and synthesized by Twist Biosciences. Sequences are listed in Data S1. The sequences were cloned into a pGeX vector containing an N-terminal smt3 tag, and a C-terminal Myc and 6xHis tag (Addgene #259391). Cloning was performed by restriction/ligation using the NEB restriction enzymes BamHI and SacII. The sequences were verified by Sanger sequencing. GAB1 was from pCDNA3.1-HA-GAB1, which was a gift from Dr. Matt Lazzara (41), cloned into the same backbone as the other phosphopeptides, and site-directed mutagenesis was performed to isolate sites Y627 and Y659, as described previously.

### Protein induction and lysis

Plasmids were transformed into BL21(DE3) E. coli cells for protein expression. An isolated colony was used to inoculate a 5 mL LB, 5 uL carbenicillin culture which grew for 16-18 hours at 37°C. 750uL of the overnight culture was added to 50mL LB media supplemented with 50ul carbenicillin and allowed to grow at 37°C to an OD600 of 0.8-1.0. The cultures were induced with 0.5 mM IPTG, and incubated at 18°C overnight. Cells were har-vested the following day by centrifugation at 8500rpm for 10 minutes, lysed via bead beating, and the lysates were stored at −70°C until purification.

### GST purification of SH2 domains

SH2 domains were purified using Pierce Glutathione Agarose (part no. 16100) following the manufacturer’s base protocol for GST purification. Beads were equilibrated in buffer (50mM Tris, 150mM NaCl) and lysate was added to the agarose for one hour to overnight, nutating at 4°C.Beads were washed three times to remove non-specifically bound proteins, with a 10-fold excess volume of equilibration buffer and centrifugation between each wash to isolate the beads. SH2 domains were then eluted with elution buffer (50 mM Tris-HCl pH 8.0, 300 mM NaCl, 500 mM imidazole). Eluted protein was dialyzed following manufacturer directions for the Pierce Slide-A-Lyzer mini dialysis with a 3.5kDa cutoff into 50mM Tris-HCl, 150mM NaCl at pH. 7.8. Proteins were aliquoted and frozen at −70°C.

### Phosphorylation of substrates

Phosphorylated substrates were generated by coexpression of the substrate with a kinase in E.coli using SISA-KiT (16). The secondary interaction used in SISA-KiT to enhance phosphorylation of the substrate is the ABL SH3 domain with the p40 polyproline sequence (APTYSPPPPP), where the tyrosine was mutated to tryptophan (W) to prevent off-target phosphorylation while still allowing for SH3 binding. We used the following kinase vectors in this study: kVh-SRC (Addgene plasmid #259387), kVh-ABL (Addgene plasmid #259379). Specific kinase/substrate pairings are listed in Data (insert table here). The substrates and kinase (Data File 1) were co-transformed into BL21(DE3) cells, and protein was expressed and co-induced with 0.5mM IPTG (substrate induction) and 0.2% L-arabinose (kinase induction) overnight at 18°C. Lysates were generated as described above, and phosphorylation of the substrates was confirmed by western blotting with a phosphotyrosine antibody (Fig. S2). Total protein content was determined using a BCA assay.

### Western Blotting

Purified SH2 domains were analyzed by SDS-PAGE to confirm protein expression. Proteins were separated on a 10% acrylamide gel and transferred onto a nitrocellulose membrane. Membranes were blocked with a 50:50 mixture of Intercept Blocking Buffer and TBS 1 hour at room temperature, followed by incubation with primary antibodies against the Myc tag for either 1 hour at room temperature or overnight at 4°C. After washing, membranes were incubated with red and green antibodies for 30 minutes at room temperature. Protein bands were visualized using a LiCor Odyssey CLx imaging system.

### Protein concentration calculation

Purified SH2 domain concentrations were calculated using a Nanodrop spectrophotometer by measuring absorbance at 280 nm. The extinction coefficient for each SH2 domain was calculated based on its amino acid sequence using the ExPASy Prot-Param tool.

### Dot blot overlay assay

Purified SH2 domains were tested for binding to phosphopeptide substrates using a modified overlay assay. Substrates were spotted onto nitrocellulose membranes in a 2-fold serial dilution using the Bio-Rad Dot Blot Apparatus (Cat: 1706545). Starting concentrations of the phosphopeptide substrates for the quadrant screens S2 were decided after determining the linear range of binding signal of the wild-type SH2 domain with each substrate in the total screen S1. The assay was performed following manufacturer’s instructions. After binding the substrates to the membrane, membranes were blocked with a 50:50 solution of Intercept Blocking Buffer and TBS for 1 hour at room temperature or 4°C overnight. Purified SH2 domains (wild type and mutants) were incubated with the membranes at a concentration of 150uM in blocking buffer for 1 hour at room temperature or 4°C overnight. After washing with TBST, membranes were incubated with anti-GST antibody (1:5000) for 1 hour at room temperature. Following three TBS washes, binding was visualized using a LiCOR Odyssey CLx imaging system.

## ACKNOWLEDGEMENTS

Research reported in this publication was supported by the National Institute Of General Medical Sciences of the National Institutes of Health under Award Numbers R35GM138127 and T32-CA-9109-46 (to Gabi Martinez). The content is solely the responsibility of the authors and does not necessarily represent the official views of the National Institutes of Health.

