## Supplementary Tables and Figures for "Studying the effect of conserved tyrosine phosphorylation within SH2 domains"

### Supplementary Materials for Martinez et al.

***This PDF file includes:.*** Table S1 to S2
Figures S1 to S9
Captions for Data S1

***Supplementary Tables.***

| Substrate | Amount ($\mu$g) | Kinase |
|---|---|---|
| PD1 Y248 | 40 | ABL1 |
| PD1 Y223 | 60 | ABL1 |
| GAB1 Y627 | 40 | SRC |
| GAB1 Y659 | 60 | SRC |
| CD33 Y340 | 40 | ABL1 |
| HGS Y216 | 40 | ABL1 |
| SIG10 Y667 | 40 | ABL1 |
| YEEI | 40 | ABL1 |
| FC$\gamma$R2B Y292 | 40 | ABL1 |
| MS4A6A Y242 | 40 | ABL1 |
| LYN Y508 | 40 | ABL1 |

**Table S1. Substrate highest protein amount used for total screens** Table of highest starting concentrations used when determining the optimal concentration for producing a linear binding signal.

| Domain | Substrate | Amount ($\mu$g) | Kinase |
|---|---|---|---|
| PTPN11 N-SH2 | PD1 Y248 | 5 | ABL1 |
| | PD1 Y223 | 30 | ABL1 |
| | GAB1 Y627 | 20 | SRC |
| | HGS Y216 | 40 | ABL1 |
| LYN SH2 | pYEEI | 15 | ABL1 |
| | MS4A6A Y242 | 20 | ABL1 |
| | LYN Y508 | 20 | ABL1 |
| | HGS Y216 | 40 | ABL1 |
| SYK C-SH2 | CD33 Y540 | 20 | ABL1 |
| | HGS Y216 | 5 | ABL1 |
| | pYEEI | 1.25 | ABL1 |
| | MS4A6A Y242 | 5 | ABL1 |

**Table S2. Substrate starting amounts for quadrant screens**. Table of substrate highest total protein amount, as determined by linear range testing done in the total phosphopeptide screens to produce a linear binding signal in the quadrant screens, by domain. Kinases used to phosphorylate the substrates are listed.

***Supplementary Figures.***

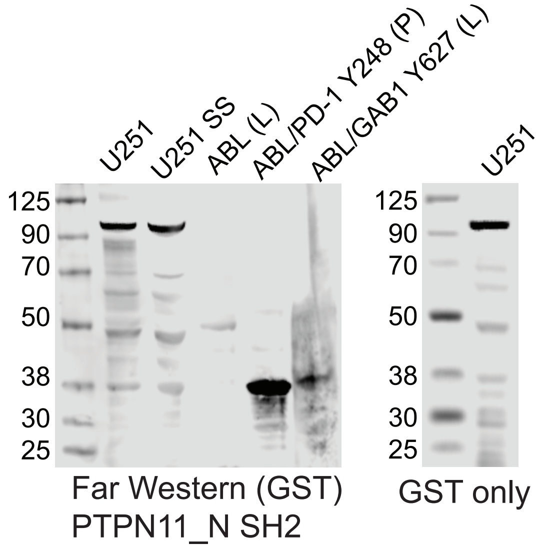


**Fig. S1. Far Western blot overlay assay using recombinant phosphoprotein substrates.** Lysates from U251 cells with and without serum starvation and recombinant phosphoproteins (PD1 pY248 and GAB1 pY627, as positive controls for SH2 domain binding) were separated by SDS-PAGE and probed with GST-fusion SH2 domains. (P) indicates the sample is purified protein, and (L) indicates that the sample was crude lysate. The lack of significant difference in the binding patterns between the GST-only and SH2 domain conditions in the U251 lysates lanes indicates that the binding observed is not largely non-specific.

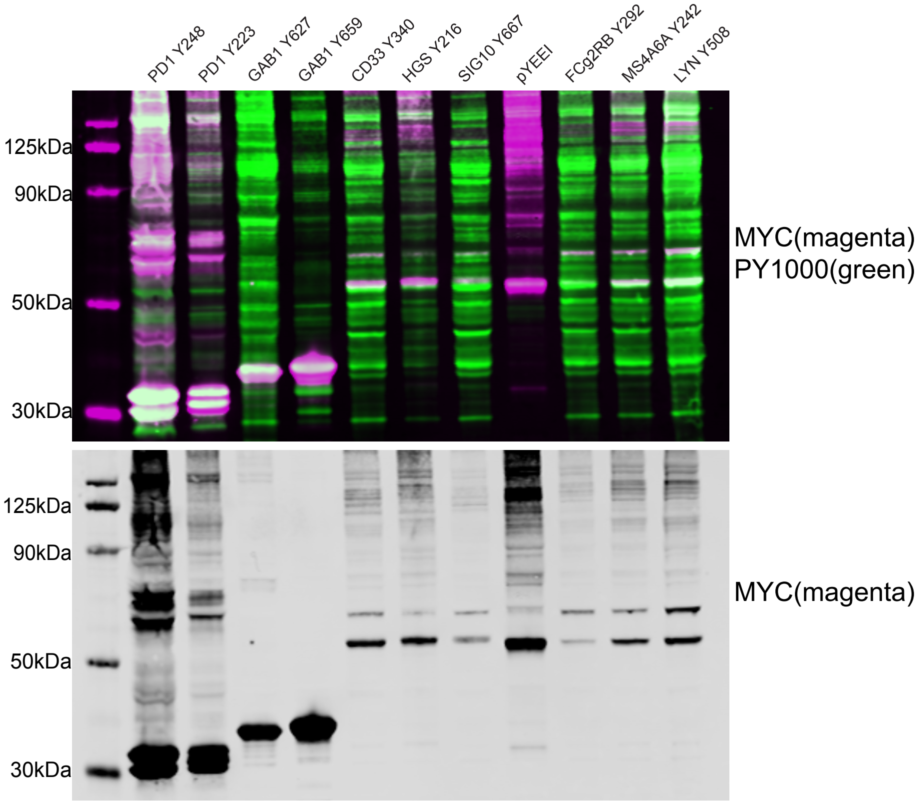


**Fig. S2. Western blot showing expression and phosphorylation of the phosphopeptide substrates used in the dot blot overlay assays.** Total protein was visualized using an anti-Myc antibody, and phosphorylation was visualized using an anti-phosphotyrosine PY1000 antibody.

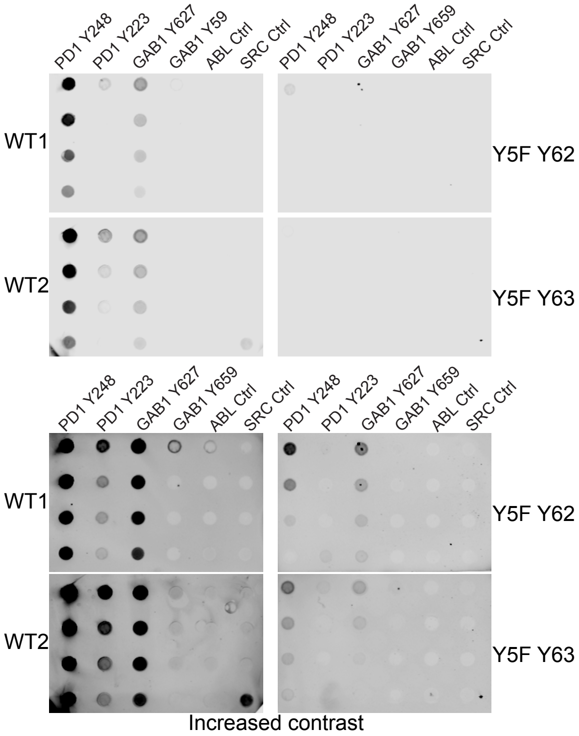


**Fig. S3. Dot Blot overlay assay results for PTPN11 N-SH2 WT comparing the ALFA WT and Y5F mutants**. Wild type were paired with Y5F Y62 and Y5F Y63, hence there are two replicates of wild type. We used an labeled ALFA nanobody for detection on all blots. The Y5F point mutation, where all but one tyrosine has been mutated to phenylalanine, has very low signal. Bottom panels are the same experiment with increased contrast to be able to determine that Y5F binding did occur, but has poor signal to noise ratio.

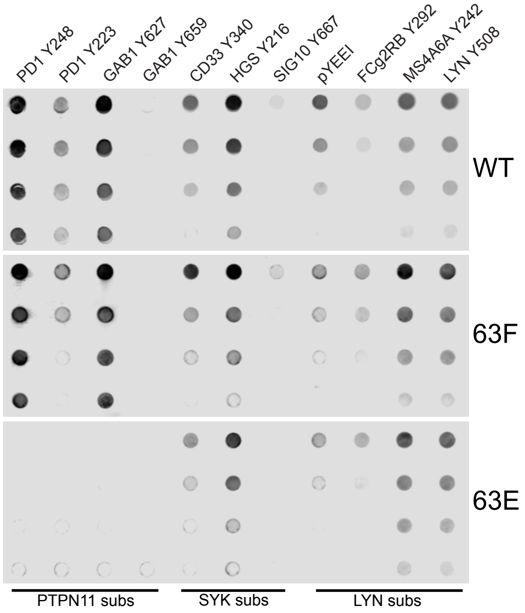


**Fig. S4. Dot blot screen analysis for PTPN11 Y63.** Dot Blot overlay assay results for PTPN11 N-SH2 WT (top), Y63F (middle), and Y63E (bottom) mutants. These SH2 domains were screened against the full panel of 11 phosphopeptide substrates.

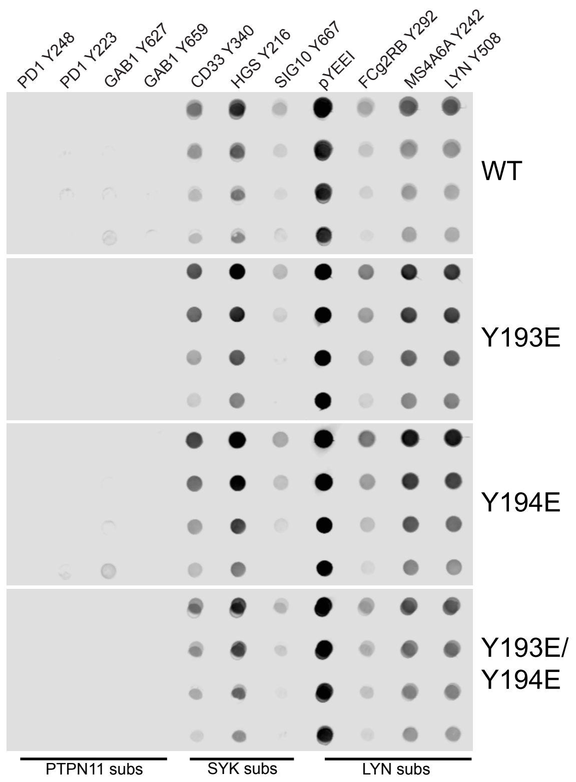


**Fig. S5. LYN Y193 and Y194 screens.** Total phosphopeptide binding screen for LYN SH2 WT, Y193E, Y194E, and Y193E/Y194E mutants.

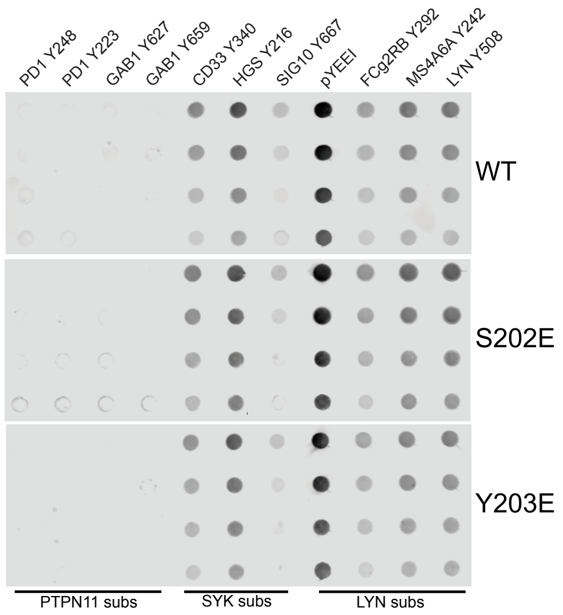


**Fig. S6. SYK S202 and Y203 screens.** Total screen binding assay results for the SYK C-SH2 WT, S202E, and Y203E mutants. 11 phosphopeptides were tested. Concentration of the substrates was diluted 2 fold down each column.

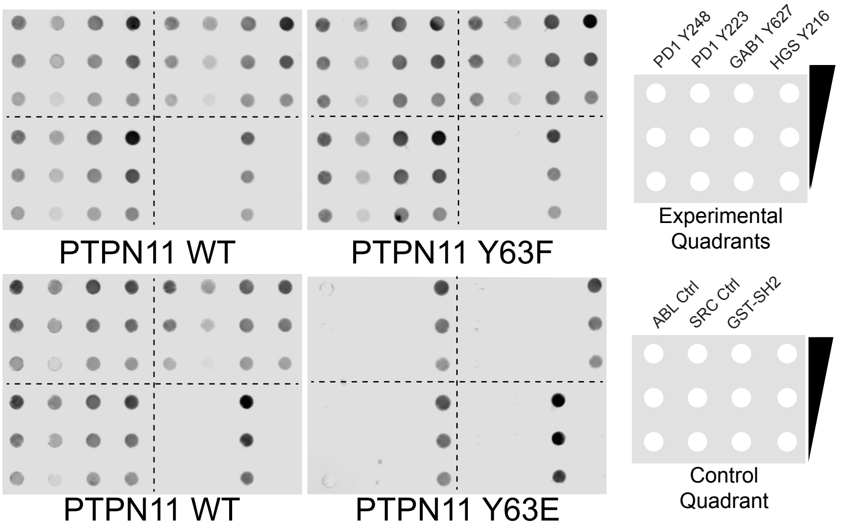


**Fig. S7. Full Membrane Results for PTPN11 N-SH2 Y63 Mutant Screening** Triplicate results for the screens on PTPN11 WT and Y63 mutants. Experimental and control membrane layouts are shown to the right of the dot blots. Membranes in the same row were imaged at the same time.

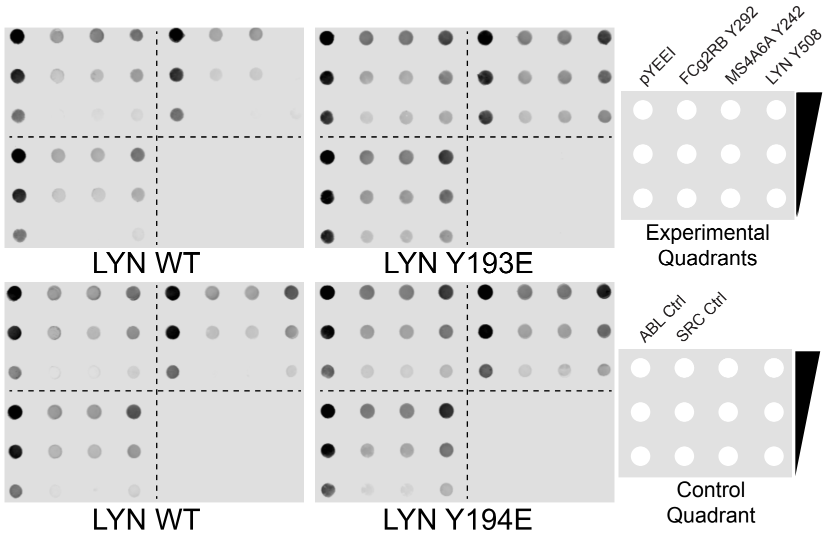


**Fig. S8. Full Membrane Results for LYN SH2 Y193E and Y194E Mutant Screening** Triplicate binding assay results for LYN SH2 WT, Y193E, and Y194E mutants. Experimental and control membrane layouts are shown to the right of the dot blots. Membranes in the same row were imaged at the same time.

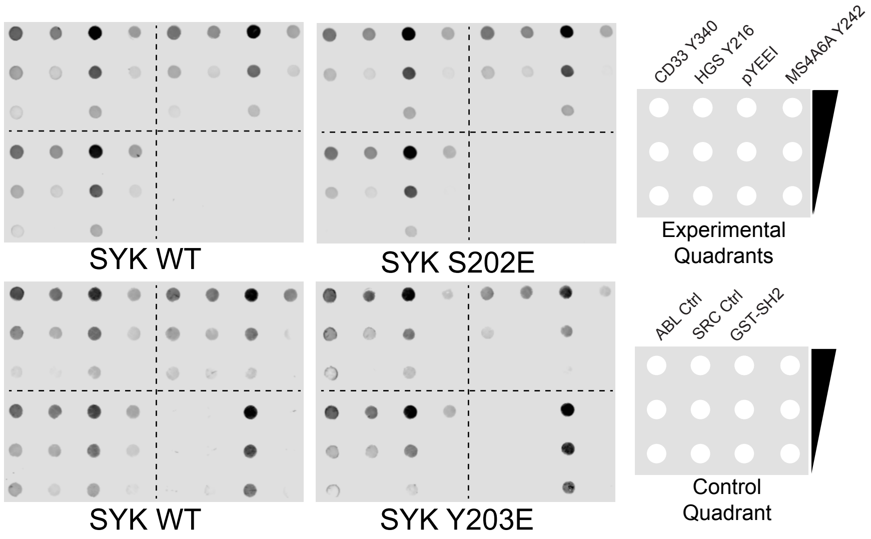


**Fig. S9. Full Membrane Results for SYK C-SH2 Y203E Mutant Screening** Triplicate binding assay results for SYK C-SH2 WT, S202E, and Y203E mutants. Experimental and control membrane layouts are shown to the right of the dot blots. Membranes in the same row were imaged at the same time.

***Supplementary Data.***

**Supplementary Data S1: Protein and DNA of SH2 domains and peptide substrates** The provided data table includes all protein record identifying information, boundaries, and sequences used for SH2 domains and peptide substrates used in this study.
